# Dynamic conformational changes of acid-sensing ion channels in different desensitizing conditions

**DOI:** 10.1101/2023.12.21.572759

**Authors:** Caroline Marcher Holm, Asli B. Topaktas, Johs Dannesboe, Stephan A. Pless, Stephanie A. Heusser

## Abstract

Acid-sensing ion channels (ASICs) are proton-gated cation channels that contribute to fast synaptic transmission and have roles in fear conditioning and nociception. Apart from activation at low pH, ASIC1a also undergoes several types of desensitization, including ‘acute desensitization’ that terminates activation, ‘steady-stated desensitization’ that occurs at sub-activating proton concentrations and limits subsequent activation, and ‘tachyphylaxis’ that results in a progressive decrease in response during a series of activations. Structural insights from a desensitized state of ASIC1 have provided great spatial detail, but dynamic insights into conformational changes in different desensitizing conditions are largely missing. Here, we use electrophysiology and voltage-clamp fluorometry to follow the functional changes of the pore along with conformational changes at several positions in the extracellular and upper transmembrane domain via cysteine-labeled fluorophores. Acute desensitization terminates activation in wild-type but introducing an N414K mutation in the β11-12 linker of mouse ASIC1a interfered with this process. The mutation also affected steady-state desensitization and led to pronounced tachyphylaxis.

Common to all types of desensitization was that the extracellular domain remained sensitive to pH and underwent pH-dependent conformational changes. These conformational changes did, however, not necessarily lead to desensitization. N414K-containing channels remained sensitive to known peptide modulators that increased steady-state desensitization, indicating that the mutation only reduced, but not precluded, desensitization. Together, this study contributes to understanding the fundamental properties of ASIC1a desensitization, emphasizing the complex interplay between the conformational changes of the ECD and the pore during channel activation and desensitization.

**Statement of significance:** Acid-sensing ion channels (ASICs) are proton-gated ion channels that contribute to synaptic activity and play roles in acidosis-related diseases. Prolonged acidosis can lead to desensitization in ASIC1a, and modulators that affect this desensitization have shown beneficial effects in pain and stroke. In this study, we investigated the functional and conformational changes during acute desensitization, steady-state desensitization, and tachyphylaxis through a mutation in the β11-12 linker of ASIC1a. We found that the mutation retained pH-dependent conformational changes of the extracellular domain (ECD) but largely disconnected these movements from the channel pore. Collectively, our work emphasizes the critical role of the β11-12 linker for the pH-dependent conformational interplay between the ECD and the channel pore.

## Introduction

Acid-sensing ion channels (ASICs) are cation-selective channels that activate upon extracellular acidification. ASICs are expressed throughout the central and peripheral nervous system, where they contribute to physiological processes such as synaptic signaling (1, 2), fear-related behaviors (3–5), and the perception and processing of pain (6, 7). Due to their involvement in acidosis-involved neuronal injury, pain, and neurological disorders, ASICs have also gained attention as possible drug targets (6, 8, 9).

In humans, four ASIC genes encode for six subunits that assemble as homo- or heterotrimers consisting of a large extracellular domain (ECD), a transmembrane domain (TMD), and intracellular C- and N-terminal domains (6, 10, 11). The ECD contains clusters of acidic residues that, when protonated during rapid drops in pH, lead to conformational changes that trigger pore opening (12, 13). The pH sensitivity of activation depends on several aspects, such as subunit composition, species, and cell type (14). In ASIC1a, the most abundant ASIC subtype in the brain, the half-maximal pH of activation (pH_50_) lies around 6.4–6.6 (14).

Besides activation, ASICs also undergo desensitization, where channels enter a non-conducting state under acidic conditions. Induction of desensitization is suggested to prevent acidosis-mediated neurotoxicity and behavioral changes (15–17). Understanding the molecular mechanisms that govern desensitization is hence of considerable therapeutic interest to e.g. achieve neuroprotection in ischemic stroke. In ASIC1a, prolonged activation with low pH leads to complete ‘acute’ desensitization, where the pore closes, and currents return to baseline within seconds (13). However, an apparent pore opening is not strictly necessary for the channels to enter a desensitized state. In ‘steady-state desensitization’ (SSD), the channels enter the desensitized state without prior opening when exposed to sub-activating proton concentrations (pH 7.3 to 6.9). Subsequent activation with low pH then leads to a reduced current amplitude (18). The degree of SSD is dependent on the proton concentration and subunit composition, but ultimately, channels need to recover from desensitization in high pH to regain the ability to activate fully (14). The different states were proposed to be either connected in a linear gating model, where a desensitized state can only be reached via the open state (14), or a branching model that allows a direct transition into a desensitized state (19, 20).

Aside from acute desensitization and SSD, ASIC1a also shows tachyphylaxis, a progressively decreased response to repeated pH stimuli (21–25). Tachyphylaxis can be observed in heterologous expression systems (e.g. (24, 26)) as well as endogenously in several cell types (21, 22, 27). Unlike acute desensitization and SSD, tachyphylaxis shows only partial and very slow recovery (25). Previous studies have suggested permeating protons, that enter the cell via ASIC1a, as key contributors to tachyphylaxis (22, 24). Factors that facilitate ASIC activation, such as low pH, small-molecule modulators, or removal of Ca^2+^ from the buffer, thus enhance tachyphylaxis (24, 25). Conversely, induction of SSD or application of a small-molecule ASIC1a inhibitor reduces tachyphylaxis (25), supporting the hypothesis that permeating protons contribute to tachyphylaxis (24). Only limited information is available about the underlying molecular mechanism, but tachyphylaxis is highly dependent on the ASIC subtype, and chimeric studies indicate a central role of residues in the N-terminal ASIC1a re-entrance loop that also affects the permeability of divalent cations (24).

From a structural perspective, the acutely desensitized states of chicken ASIC1 represent an intermediate between the resting and open states (12, 13, 28). The upper half of the ECD of the desensitized state is structurally highly similar to that of the open state. Both feature a collapsed acidic pocket (an ECD region that houses a cluster of protonatable side chains) in response to the acidic conditions under which they were resolved. On the other hand, the lower ECD and TMD of the desensitized states are structurally more similar to those of the resting state, featuring a closed pore (12, 13, 28). The transition zone between the upper and lower ECD includes the β11-12 linker with residues L414 and N415 showing substantial rearrangement between the open and desensitized state (Figure 1A). Crosslinking studies, simulations, and several functional analyses of mutations of residues L414, N415, and in their vicinity, Q277 (chicken ASIC1 numbering) have emphasized the importance of this linker for activation and acute desensitization (20, 29–33). Therefore, the region is often referred to as ‘molecular clutch’ (12). When engaged, the clutch transmits the pH-dependent conformational changes of the upper ECD to the TMD to open the pore. However, when the β11-12 linker switches conformation, as seen in the acutely desensitized state, the clutch is disengaged, leading to a decoupling between the ECD and the pore. While these insights offer a compelling molecular mechanism of acute desensitization, less is known about the ECD conformational changes responding to a more gradual decrease in pH during SSD or the responsiveness of the ECD during tachyphylaxis.

**Figure 1:**
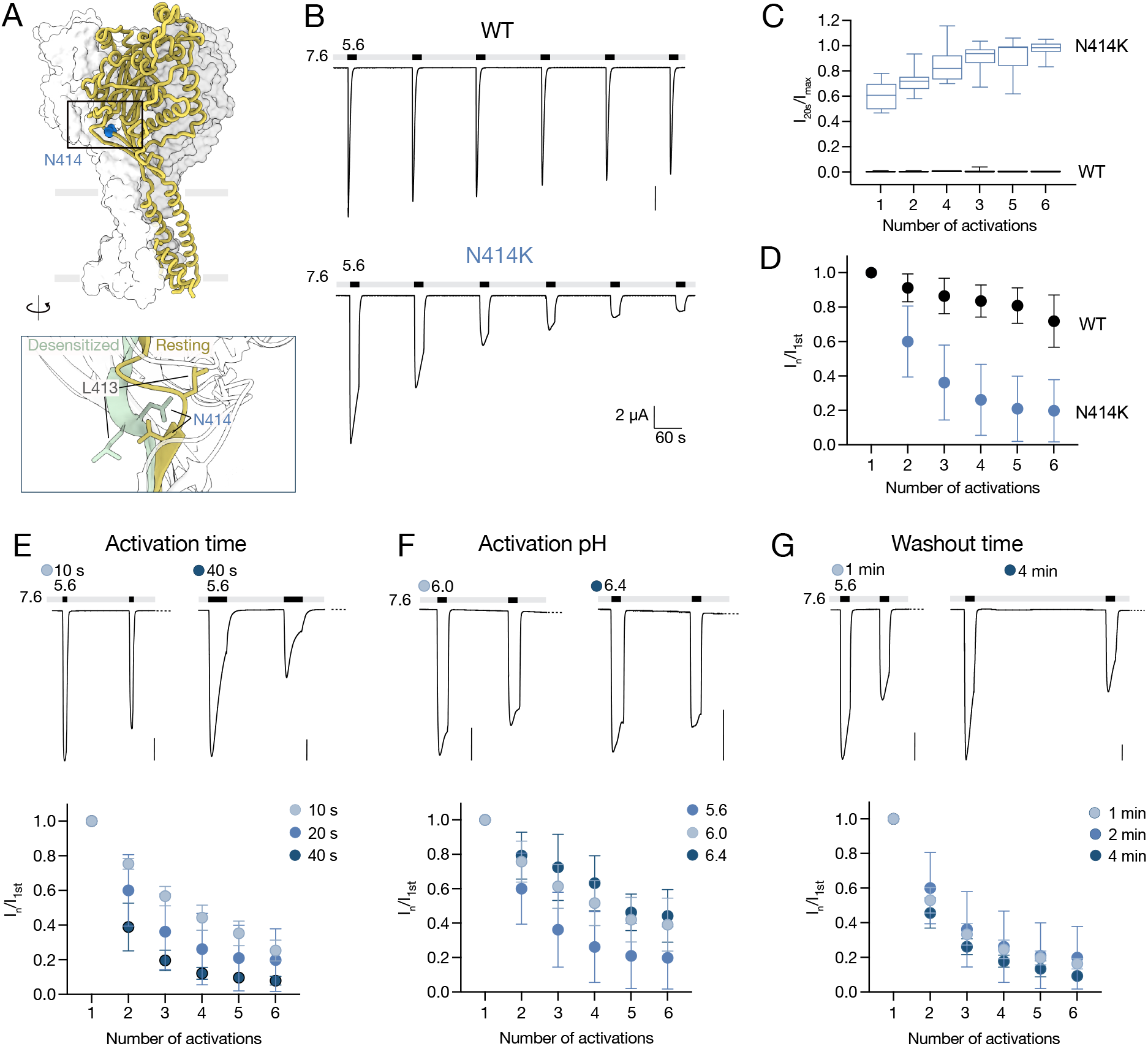
Introduction of the N414K mutation in mASIC1a affects desensitization. **A**) Structural overview of a resting state of chicken ASIC1 (PDB ID 6VTL) with position corresponding to mASIC1a N414 highlighted in blue. Inset shows a closeup of how residues L413 and N414 in the β11-12 linker change position between a resting and a desensitized (PDB ID 6VTK, pale green) state. **B**) Representative TEVC trace of mASIC1a WT (top) and N414K (boVom) showing repeated 20 s activation with pH 5.6, with 2 min recovery time in pH 7.6. **C**) Assessment of acute desensitization by comparing currents at the end of the 20 s activation to the maximal current in the beginning of the peak (n=8). **D**) Assessment of tachyphylaxis by comparing currents of recurrent activations to the current of the first activation (n=8-12). **E**) Top: Section of a TEVC trace showing the first two out of six activations to show the effect of activation time on acute desensitization and tachyphylaxis of mASIC1a N414K. BoVom: Quantitative analysis of tachyphylaxis data normalizing the current of each activation to the first (n=4-5). **F**) Same as in panel E, but using 20 s activation of varying pH (n=4-5).**G**) Same as in panel E, but the 20 s activations at pH 5.6 were intercepted by various washout times (n=3-8). All scale bars are 2 μA and traces are displayed on the same time scale. Data in graphs of panels D-G are presented as mean ± SD.

A better understanding of the ASIC1a desensitization mechanisms is important since several known ASIC1 modulators, such as the endogenous neuropeptide Big dynorphin and many animal-derived toxins, affect desensitization (16, 34). We therefore set out to investigate how the ECD responds to the different types of desensitization and how they influence each other. We used the N414K mutation in the β11-12 linker of mouse ASIC1a (mASIC1a), which is known to dramatically affect acute desensitization, to assess its effect on SSD and tachyphylaxis using an approach that allows us to simultaneously measure channel function and ECD conformational changes. We observed that the ECD and upper TMD underwent pH-dependent conformational changes during all types of desensitization but that these movements did not always translate into functional changes at the level of the pore. Finally, we demonstrated that peptide modulators known to affect ASIC1a desensitization still interact with channels even when desensitization is disrupted.

## Methods

### Molecular biology

Mouse ASIC1a cDNA in the pSP64 vector was obtained from Marcelo Carattino (University of Pittsburgh). The rat ASIC3 clone in the pRSSP6009 vector was provided by Stefan Gründer ( RWTH Aachen University). Mutations were introduced via site-directed mutagenesis using PfuUltraII Fusion polymerase (Agilent) and custom-made DNA mutagenesis primers (Eurofins Genomics). Mutagenesis was confirmed by sequencing the full coding frame (Eurofins Genomics or Macrogen). All cDNAs were linearized using EcoRI. For transcription to capped cRNA, the Ambion mMESSAGE mMACHINE SP6 kit (Thermo Fisher Scientific) was used.

### Electrophysiological setups

Oocytes from female *Xenopus laevis* frogs were surgically removed and prepared as previously described (35). Isolated oocytes were injected with 0.5–50 ng of cRNA (volumes between 20 and 50 nL) and incubated for 2–5 days at 18°C in supplemented ND96 solution, containing 96 mM NaCl, 2 mM KCl, 1.8 mM CaCl_2_, 1 mM MgCl_2_, 5 mM HEPES along with 2.5 mM sodium pyruvate, 0.5 mM theophylline, 0.05mg/ml gentamycin and 0.05 mg/ml tetracycline, adjusted to pH 7.6. Before conducting VCF experiments, oocytes were labeled by placing them for 30 minutes in OR2 solution containing 82.5 mM NaCl, 2 KCl mM, 1 MgCl_2_ mM, pH 7.4 along with 10-30 μM Alexa Fluor 488 C5-maleimide (Thermo Fisher Scientific). After labeling, oocytes were washed twice with OR2 solution and stored in the dark until further use.

The recording setup was the same as described previously (36). Oocytes were placed in a recording chamber (37) and continuously perfused with Ca^2+^-free ND96 (96 mM NaCl, 2 mM KCl, 1.8 mM BaCl_2_, 1 mM MgCl_2_, 5 mM HEPES) adjusted to the desired pH value with NaOH or HCl. In experiments where we use PcTx1 or Big dynorphin, buffers were supplemented with 0.05% bovine serum albumin (≥98% essentially fatty acid-free, Sigma-Aldrich). Solutions were exchanged using a gravity-driven 8-line automated perfusion system operated by a ValveBank module (AutoMate Scientific). VCF recordings were done using an inverse microscope (IX73 with LUMPlanFL N ×40, Olympus), a halogen lamp as the light source (Olympus TH4-200), and a standard GFP filter set (Olympus). The emission was detected using a P25PC-19 photomultiplier tube (Sens-Tech) and photon-to-voltage converter (IonOptix). Microelectrodes (borosilicate capillaries 1.2 mm OD, 0.94 mm ID, Harvard Apparatus) were backfilled with 3 M KCl, leading to resistances of 0.3–1 MΩ. An OC-725C amplifier (Warner Instruments) was used to acquire signals at 1000 Hz, filtered by a 50–60 Hz noise eliminator (Hum Bug, Quest Scientific). Signals were digitized using an Axon Digidata 1550 Data Acquisition System and recorded with pClamp software (10.5.1.0, Molecular Devices). Prior to analysis, current signals were further digitally filtered at 3 Hz using an 8-pole Bessel low-pass filter. Displayed current traces have been subjected to an additional 10× data reduction, unless stated otherwise.

PcTx1 (>95% purity) was obtained from Alomone Labs. Big dynorphin was synthesized using automated solid-phase peptide synthesis and purification via reversed-phase high-performance liquid chromatography as described previously (36). Peptide stock solutions were prepared in MilliQ water (18.2 MΩ resistivity) and stored at –20°C. Prior to recording, the peptides were diluted to the desired concentration in BSA-containing buffer.

### Electrophysiological recording protocols

Recordings were conducted at -40 mV, except for data in Suppl. Figure S2F and G, where oocytes were clamped at -10 mV. For tachyphylaxis experiments, ASIC-expressing oocytes were exposed to pH 5.6 for 20 s, followed by a washout in pH 7.6 for 60 s, unless stated otherwise. A running buffer at pH 7.6 was used for concentration-response relationships of current and fluorescence. Channels were activated for 20 s at pH 5.6, washed for 20-60 s with buffer at pH 7.6, preconditioned for 60 s (9 min for data in Suppl. Figure S2F) at moderate pH, and then directly activated. This protocol was repeated with varying preconditioning pHs, resulting in pH 5.6 control activations between each SSD measurement, enabling us to account for tachyphylaxis. In TEVC experiments with PcTx1, 30 nM PcTx1 was applied for 120 s at pH 7.4 for WT or at a concentration of 100 nM in pH 6.7 for K105C/N414K, followed directly by a short activation at low pH. For VCF experiments, 300 nM PcTx1 was used at pH 7.4. Big dynorphin was applied at 1 μM in pH 7.6 in all experiments following the same protocol used for PcTx1, except that it was applied at pH 6.6 in the K105C/N414K mutant.

### Data analysis

Data analyses were performed in Prism (10; GraphPad Software). The fluorescence baseline was adjusted for all fluorescence traces before analysis. For tachyphylaxis experiments, peak currents and peak fluorescence signals of each activation were normalized to the response of the first activation. For Figure 1D, current amplitudes at the end of the 20 s pH 5.6 activation were normalized to the peak current of the same activation. For pH-response data, peak currents were normalized to the average of the flanking low-pH activations, and fluorescence amplitudes were normalized to the neighboring low-pH response. Data was plotted as mean responses ± SD for each tested pH. Where curves are indicated, normalized pH responses were fitted with a Hill equation constrained at min=0 (for the fluorescence in Figure 4F, no constraints were applied) and data was reported for averaged pH_50_ values and upaired Welch’s t-tests were used for comparisons. For display in figures, a single fit to the average normalized responses is shown. For PcTx1 and Big dynorphin experiments, current and fluorescence responses were normalized to the flanking/neighboring low-pH activations as described for SSD measurements. A paired t-test was used to compare data with and without peptides. For comparison of the fluorescence intensity in Suppl. Figure S4C, an ordinary one-way ANOVA with a Dunnett’s multiple comparisons test was used. Fluorescence intensity was calculated as ‘max. fluorescence’/’ baseline fluorescence’ in percent.

All data points are presented as mean ± SD unless stated otherwise, and the number of replicates (n) represents individual experimental oocytes. Results were obtained from at least two batches of oocytes. All graphs and illustrations were made in Prism (10.0, GraphPad Software) and AffinityDesigner, and structure rendering was done using ChimeraX (38). Sequence alignment was done using Clustal Omega (39).

## Results

### The N414K mutation reduces acute desensitization and enhances tachyphylaxis

The β11-12 linker of ASIC1 is a critical structural region underlying desensitization. Mutations of key residues in this linker including L415 and N416 in human ASIC1a (29) or the corresponding residues in chicken ASIC1 (31) reduce the rate of acute desensitization and shift SSD to more acidic pH values. We mutated mASIC1a residue N414 to lysine (Figure 1A), expressed it in *Xenopus laevis* oocytes, and recorded the response to low pH stimuli using two-electrode voltage clamp (TEVC) electrophysiology. When activated with pH 5.6 interspersed with 2-minute washout intervals, the acute desensitization of the N414K mutant was markedly reduced compared to wild-type (WT) (Figure 1B). While the WT current completely desensitized within the 20 s exposure to pH 5.6, 60% of the maximal currents remained for the N414K mutant even at the end of the 20 s application (Figure 1C). The recordings also revealed changes in peak shape over the course of recurrent activations; acute desensitization was generally more pronounced in the first activation and decreased over recurrent activations (Figure 1B, C). Besides a decrease in the extent of acute desensitization, the N414K mutant also showed pronounced tachyphylaxis, as seen by the reduction in current amplitude upon recurrent activation with pH 5.6 (Figure 1B and D). Note that the size of a current is similar to the values reached at the end of the preceding activation. The extent of tachyphylaxis was more pronounced when activation times were longer (40 s) and smaller when the activation times were shorter (10s) (Figure 1E). Tachyphylaxis also depended on the pH used for activation. For example, activation with pH 6.4 and 6.0 led to less tachyphylaxis than when pH 5.6 was used for activation (Figure 1F). By contrast, longer (4 min) or shorter (1 min) washout times had little effect on the extent of tachyphylaxis (Figure 1G). Lower pH and longer activation time can increase the permeation of protons and Ca^2+^ that can flow through ASIC1a together with Na^+^ (22, 24). Therefore, our results align with the hypothesis that these permeating ions govern tachyphylaxis (24).

The ASIC3 isoform shows faster acute desensitization than ASIC1a and exhibits no canonical tachyphylaxis (24). To understand whether the absence of tachyphylaxis in ASIC3 is due to the short openings or an underlying structural difference, we introduced the N414K mutation to rat ASIC3 (rASIC3). The mutation strongly reduced acute desensitization but only introduced around 40% of tachyphylaxis after six consecutive activations, even when channels were activated for 40 s. (Suppl. Figure S1A and B). These results indicate that the propensity for tachyphylaxis comes from inherent differences between mASIC1a and rASIC3. Since the intracellular N-terminal domain was suggested to be involved in the mechanism that governs tachyphylaxis along with permeating protons (24), we mutated some of the negatively charged side chains of the mASIC1a N-terminal domain to their counterparts found in rASIC3 (Suppl. Figure S1C). However, we did not see any significant effect of these mutants on the development of tachyphylaxis (Suppl. Figure S1D), warranting a more in-depth study in the future.

Taken together, introduction of the β11-12 linker mutation N414K led to reduced acute desensitization and enhanced tachyphylaxis, with tachyphylaxis being dependent on the duration and pH of activation.

### In the N414K mutant, upper- and mid-ECD respond to moderate pH, but SSD is reduced

We set out to better understand the ECD conformational changes that accompany the different types of desensitization and examine the impact of the N414K mutant on the communication between the ECD and the pore. To this end, we introduced an environmentally sensitive fluorophore to the ECD, to then simultaneously followed changes in fluorescence and current in VCF experiments. To attach the fluorophore, we replaced residue K105 with a cysteine (K105C, Figure 2A) and labeled it with Alexa Fluor 488 C_5_ maleimide. VCF readouts from this position have shown to act as a proxy of ECD conformational changes during mASIC1a gating and peptide binding (36, 40, 41). Since channel kinetics exceed our perfusion speed, we focus our observations on conformational changes related to functional states rather than gating transitions. Consistent with previous work (40, 41), we show that the Alexa Fluor 488 C_5_ maleimide-labeled K105C mutant responded to pH 5.6 activation with an inward current that acutely desensitizes within the 20 s application frame, accompanied by an upward deflection of the fluorescent signal (Figure 2B, left, Suppl. Figure S2A). The fluorescent signal typically displayed a transient peak that settled into a plateau, likely reflecting channel opening and desensitization, respectively (41). During six consecutive activations with pH 5.6, the K105C mutant showed minor tachyphylaxis with near-identical fluorescence signals (Figure 2C, left). The introduction of N414K into the K105C mutant (K105C/N414K) showed a similar fluorescence response as seen in K105C (Figure 2B, right, Suppl. Figure S2A). Like the N414K single mutant, the K105C/N414K double mutant showed pronounced tachyphylaxis of the current (Figure 2B, right). Again, the extent was dependent on activation time and pH, but not washout time (Suppl. Figure S2B-D). However, while the current decreased over recurrent activations, the level of fluorescence remained stable throughout the activation protocol, indicating that the top of the ECD remains conformationally responsive to pH (Figure 2B and C, left).

**Figure 2:**
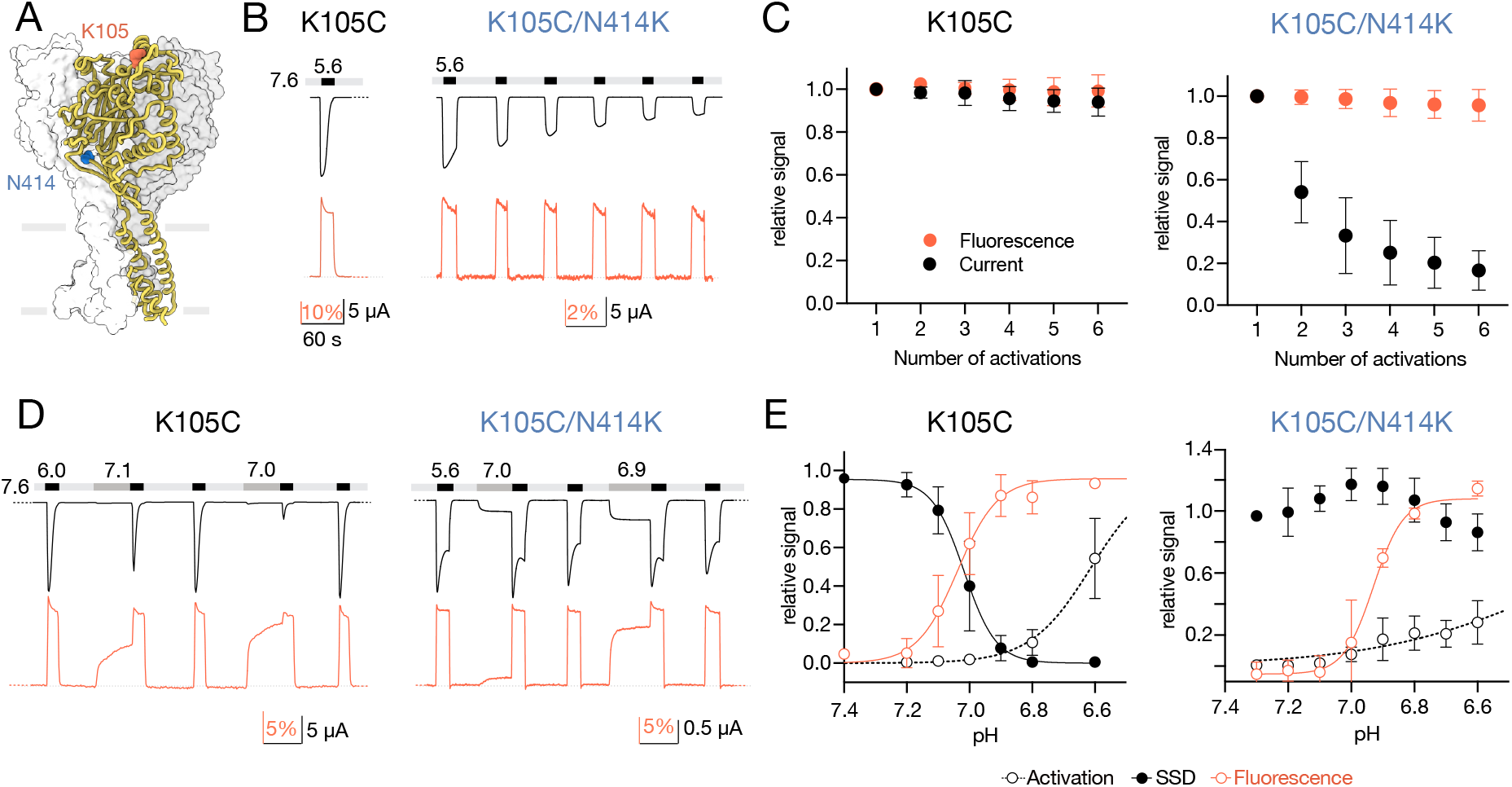
The N414K mutant undergoes ECD conformational changes but shows reduced propensity for SSD. **A**) Structural overview of cASIC1 (PDB ID 6VTL) showing positions corresponding to mASIC1a N414 in blue, and K105 used for fluorescent labeling in red. **B**) Representative VCF trace of mASIC1a K105C and K105C/N414K labelled with Alexa Fluor 488, with the current in black and the fluorescence in red. The trace for K105C/N414K shows repeated 20 s activation with pH 5.6 with 1 min recovery time in pH 7.6. **C**) Assessment of tachyphylaxis of K105C (leg) and K105C/N414K (right) by comparing currents and fluorescence of recurrent activations to values of the first activation (n=5-6). **D**) Representative excerpts of VCF recordings of K105C (leg) and K105C/N414K (right) to assess the pH sensitivity of the fluorescent signal (red) and SSD (current, black). For this purpose, channels were preconditioned for 60 s in moderate pH (dark gray bars) before they were activated for 20 s at low pH (black bar). Each such measurement was flanked by activation with low pH to account for tachyphylaxis. **E**) Quantitative analysis of recordings in panel D. Currents at moderate pH (=activation), currents evoked ager preconditioning in moderate pH (=SSD), and fluorescence were normalized to the average of the flanking currents at low pH (n=3-9). Traces in panels B and D are all on the same time scale. Data in graphs are presented as mean ± SD.

We then wanted to further assess the responsiveness of the ECD to pH and investigate if the decrease in acute desensitization also led to diminished SSD. We thus preconditioned the channels for 60 s in moderate pH (7.4-6.8) before activating them with low pH. For the K105C mutant, this protocol induced SSD in the preconditioning pH range of 7.2 to 6.8 (pH_50_SSD_ 7.04 ± 0.06, Figure 2D and E, left, Suppl. Table T1). Simultaneously, the fluorescence signal increased within the same pH range as SSD (pH_50_Fluorescence_ 7.06 ± 0.06, Suppl. Table T1), indicating that the pH-induced conformational changes detected by the fluorophore are indeed related to SSD, as indicated previously (36). Since tachyphylaxis in the N414K mutant could lead to a biased observation of SSD due to decreasing current even without preconditioning, we recorded low pH responses between each SSD measurement. We then normalized the preconditioned current to the mean size of the two flanking low pH activations (Figure 2D, Suppl. Figure S2E). When accounting for tachyphylaxis, we did not observe typical SSD for the K105C/N414K mutant within pH 7.3 to 6.5 (Figure 2D and E, right). Activation and tachyphylaxis make it challenging to extend the pH range towards lower pH, however, within this pH range, the ECD underwent pH-dependent conformational changes similar to those observed in the K105C mutant (pH_50_Fluorescence_ 6.93 ± 0.02, Figure 2D and E, Suppl. Figure S2E, Suppl. Table T1). These conformational changes thus even appear when efficient SSD is impeded but seem also disconnected from the relatively shallow pH-dependent activation of the K105C/N414K mutant (pH_50_Activation_ 6.49 ± 0.25, Suppl. Table T1). Limited SSD in the pH range tested could originate from non-equilibrium conditions during the preconditioning step, especially given the slow entry into acute desensitization of the K105C/N414K mutant. We thus preconditioned K105C/N414K in pH 6.8 for 9 min (Suppl. Figure 2F). Although the subsequent pH 5.6 current was indeed slightly smaller than when preconditioned for 1 min (relative currents 0.89 ± 0.15 (9 min), 1.09 ± 0.15 (1 min,) Suppl. Figure S2G, Suppl. Table T2), SSD remained marginal compared to the almost maximal fluorescence response (relative response 0.98 ± 0.04). While we cannot rule out that the K105C/N414K mutant can fully steady-state desensitize at lower preconditioning pH and/or longer application time, our data highlights how the conformational changes in response to application of moderate pH are mostly disconnected from functional outcomes.

The insights align with the notion that the N414K mutation impedes isomerization of the β11-12 linker and thus the onset of acute and SSD.

Besides the β11-12 linker, previous studies have also identified the neighboring β1-2 linker to be of importance for activation and desensitization (12, 33, 42–44). We therefore introduced a cysteine mutation at mASIC1a residue V80, located at the beginning of the β1-2 linker (V80C, Figure 3A), to fluorescently label it as previously described (40, 41). Mutation and labeling of V80C led to currents that promptly activated and desensitized upon activation with pH 5.6, similar to WT (Figure 3B, left, Suppl. Figure S3A). The fluorescence signal that accompanied the current was negative in amplitude but had a similar shape to that of K105C, with a sharp initial peak followed by a plateau. When combined with the N414K mutation (V80C/N414K), acute desensitization was strongly reduced, resulting in only partial desensitization within a 20 s activation with pH 5.6, and similar fluorescence response compared to the V80C single mutation (Figure 3B, right, Suppl. Figure S3B). Consecutive activations with pH 5.6 evoked strong tachyphylaxis in V80C/N414K, but not in V80C, while the fluorescence response remained stable across six consecutive activations for both mutants (Figure 3B, right, and 3C). In V80C, the changes in fluorescence appeared in a similar pH range as SSD (pH_50_Fluorescence_ 7.36 ± 0.04, pH_50_SSD_ 7.17 ± 0.03), while activation happened at lower pHs (pH_50_Activation_ 6.65 ± 0.22, Figure 3D and E, left, Suppl. Figure S3C, Suppl. Table T1), as reported earlier (41). On the other hand, SSD of the V80C/N414K mutant showed only a shallow pH dependence and was not fully resolved within the pH range tested (Figure 3D, and E, right, Suppl. Figure S3D). The pH-dependence of the conformational changes reported by the fluorophore at V80C (pH_50_Fluorescence_ 7.11 ± 0.03, Suppl. Table T1) thus neither appear to report on SSD nor activation.

**Figure 3:**
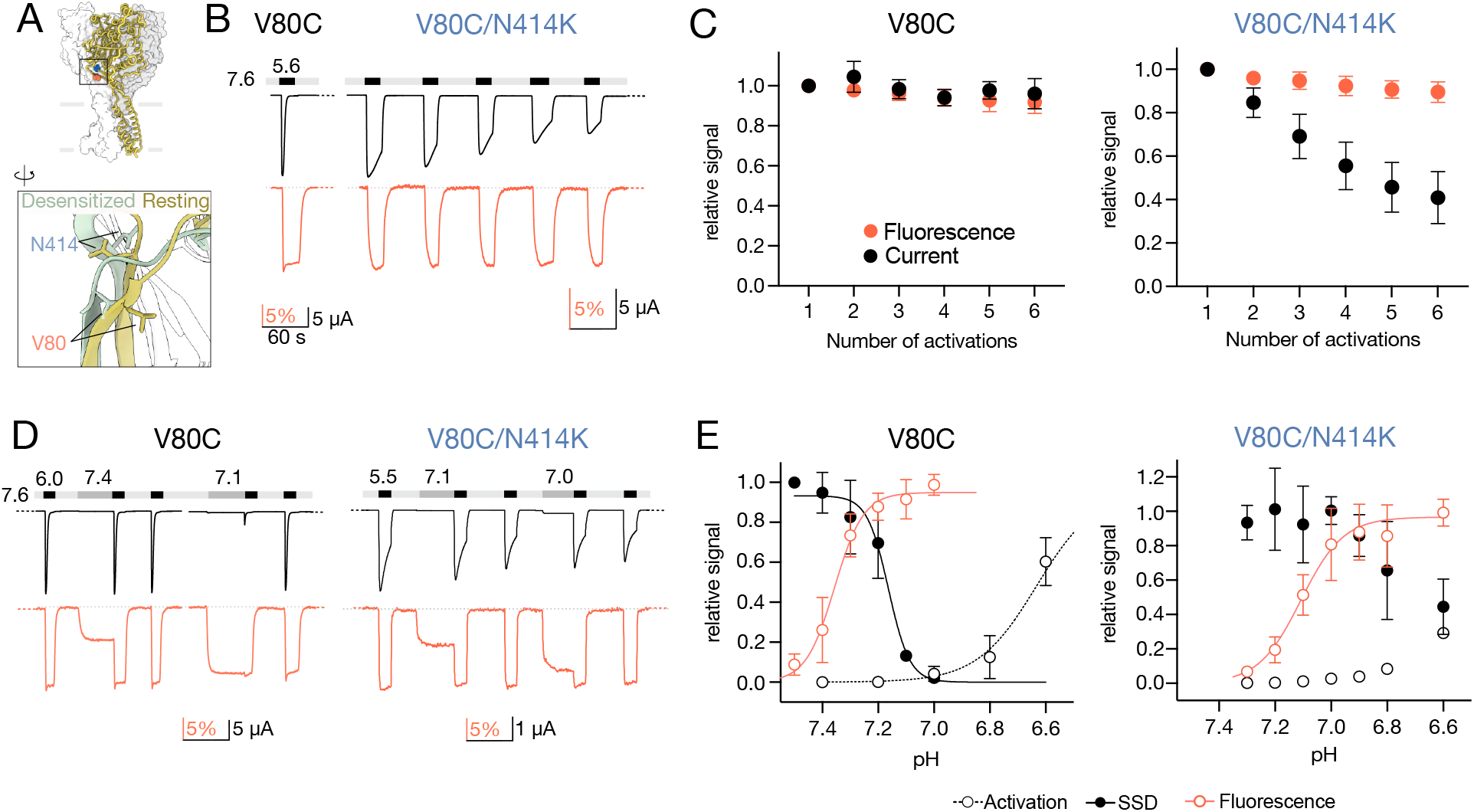
The N414K mutation appears to impede conformational changes of the mid-ECD to elicit desensitization. **A)** Structural overview and zoom of cASIC1 (resting, yellow; desensitized, pale green) showing positions corresponding to mASIC1a N414, and V80 used for fluorescent labeling. **B**) Sections of VCF traces of mASIC1a V80C (leg) and V80C/N414K (right) with the current in black and the fluorescence in red. The trace for V80C/N414K shows repeated 20 s activation with pH 5.6 with 1 min recovery time in pH 7.6. **C**) Assessment of tachyphylaxis of V80C (leg) and V80C/N414K (right) comparing currents and fluorescence of each activation to the first activation (n=4-5). **D)** Representative excerpts of VCF recordings of V80C (leg) and V80C/N414K (right) to assess the pH sensitivity of the fluorescent signal (red) and SSD (current, black) as described for K105C. **E)** Fluorescence, activation, and SSD of V80C (leg) and V80C/N414K (right) assessed as described in Figure 2E (n=3-12). Traces in B and D are displayed on the same time scale. Data in graphs are presented as mean ± SD.

Overall, labeling at V80C had comparable outcomes as labeling in K105C, indicating that even at the structural transition zone of the upper and the lower ECD, fluorophore labeling reports on ECD conformational changes that are not efficiently translated to desensitization in the N414K mutant background.

### Labeling in the upper TMD domain reports on activation and desensitization

We went on to examine if the conformational changes in the upper TMD would resemble the ones seen for the upper ECD or if they instead follow the functional changes of the TMD. To this end, we labeled the channel in two more positions previously reported to yield changes in fluorescence signals, E425C and H72C (Figure 4A, Suppl. Figure S4A) (19, 40, 45).

**Figure 4:**
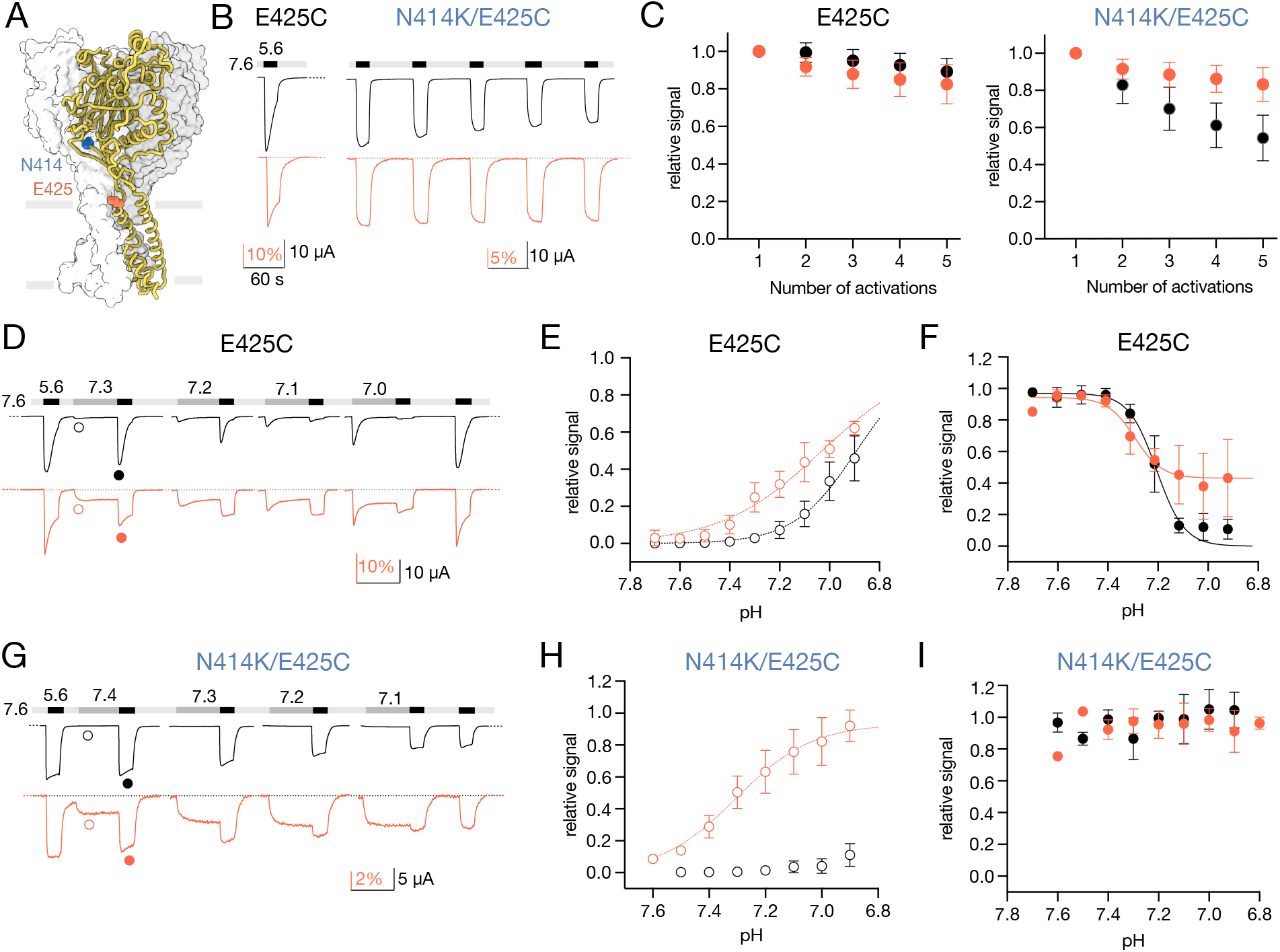
E425C reports on conformational changes related to activation and SS.D. **A)** Structural overview of cASIC1 (PDB ID 6VTL) showing positions corresponding to mASIC1a N414 in blue, and E425 used for fluorescent labeling in red. **B)** Representative VCF trace of mASIC1a E425C and N414K/E425C labeled with Alexa Fluor 488, with the current in black and the fluorescence in red. The trace for N414K/E425C shows repeated 20 s activation with pH 5.6 with 1 min recovery in pH 7.6. **C)** Assessment of tachyphylaxis of E425C (leg) and N414K/E425C (right) by comparing currents and fluorescence of recurrent activations to values of the first activation (n=16-22). **D)** Representative excerpt of VCF recordings of E425C to assess the pH sensitivity of the fluorescent signal (red) and SSD (current, black). For this purpose, channels were preconditioned for 60 s at moderate pH (dark gray bars) before activating for 20 s at low pH (black bar). **E)** Quantitative analysis of current and fluorescence evoked by moderate pH, marked as an example in panel D with open circles (n=4-19). **F)** Quantitative analysis of current and fluorescence evoked by low pH ager preconditioning in moderate pH. Marked as an example in panel D with closed circles (n=4-19). **G-I)** Same as in D-F, respectively, but for N414K/E425C (n=3-19). All traces are on the same time scale. Data in graphs are presented as mean ± SD.

Residue E425C is located at the extracellular end of TM2, and mutation and labeling of this position reduced acute desensitization of the channel but led to robust fluorescence signals (Figure 4A and B, left, Suppl. Figure S4B, top). Despite the slower acute desensitization, the E425C mutant only showed slight tachyphylaxis, which was accompanied by a small stepwise fluorescence decrease upon recurrent activation (Figure 4C, left). Introduction of the N414K mutation on the E425C background (N414K/E425C) led to channels that showed only a minor degree of acute desensitization and a fluorescent signal that slowly reached a plateau during acute desensitization (Figure 4B, right, Suppl. Figure S4B, bottom). Tachyphylaxis was robust (Figure 4B and C, right), although markedly smaller in its extent than seen for K105C/N414K (Figure 2B and C) or V80C/N414K (Figure 3B and C). Meanwhile, the fluorescence signal amplitude decreased slightly with repeated activations, similar to what we observed with the E425C single mutant (Figure 4B and C).

To investigate if the slight decrease in fluorescence signal could reflect tachyphylaxis, we used the same SSD protocol as described for the previous two labeling positions (Figure 4D). Notably, the fluorescence signal of the E425C mutant shallowly increased from pH 7.5 to 6.8 and was only slightly right-shifted from the pH-dependent activation (pH_50_Fluorescence_ 7.06 ± 0.07, pH_50_Activation_ 6.87 ± 0.10, Figure 4E, Suppl. Table T1). SSD remained intact with a pH_50_SSD_ of 7.17 ± 0.13 (Figure 4F, Suppl. Table T1). Interestingly, low pH activation after preconditioning in moderately low pH led to a decreased fluorescence signal (pH_50_Fluorescence_SSD_ 7.29 ± 0.04), roughly mirroring the SSD of the current (Figure 4F, Suppl Table T1). These observations indicate that labeling in position E425C indeed seems to report on conformational changes of the TM2 (or its vicinity) that are related to pore opening. Notably, the pH range of activation and the pH-dependence of the fluorescence signal did not completely overlap (Figure 4E), and the reduction of the fluorescence after preconditioning was only partial despite almost complete SSD in this range (Figure 4F). These insights indicate that the fluorescence signal not only reports on pore opening but is likely convoluted from conformational changes related to desensitization, similar to previous observations (45). These findings are thus distinct from those obtained from the upper and mid-ECD, where the pH-dependence of the fluorescence appeared to mostly follow desensitization. Introduction of the N414K mutation yielded channels that were activated at a lower pH and showed no SSD within the pH range tested (Figure 4G-I). In contrast, the mutation had little effect on the pH dependence of the fluorescent signal (pH_50_Fluorescence_ 7.22 ± 0.15, Suppl. Table T1).

To probe if labeling the upper end of TM1 would lead to similar results, we turned to H72C (Suppl. Figure S4A). While the fluorescence signals thus far have been robust, labeling at position H72 only led to small fluorescence signals comparable in size to those recorded, when WT-expressing *Xenopus* oocytes were exposed to Alexa Fluor 488 dye (Suppl. Figure S4C). A likely culprit seemed to be residue C70, which usually does not appear to label efficiently or at least yield no significant fluorescence change (Suppl. Figure S4C). Introduction of a neighboring cysteine, H72C, could however lead to disulfide bond formation and thus limit labeling. We therefore generated the C70S mutant, which significantly increased the fluorescence signals in the C70S/H72C mutant (Suppl. Figure S4C) and resulted in channels that desensitized within 20s (Suppl. Figure S4D and E). This double mutant showed a minor degree of tachyphylaxis, accompanied by a fluorescence signal that also slightly decreased over repeated activations (Suppl. Figure S4F, left). The pH dependence of the fluorescence was reminiscent of that seen for E425C (Figure 4E and F), but the inherent variability in the data obtained from this mutant made it difficult to definitively answer if the fluorescence signal recorded at this position also reports on both pore opening and ECD protonation (Suppl. Figure S4G). As expected, additional introduction of N414K (C70S/H72C/N414K) decreased acute desensitization and SSD but enhanced tachyphylaxis (Suppl. Figure S4D-F). The pH dependence of the fluorescence signal appeared similar to the double mutant but did not seem to occur at the same pH range as SSD (Suppl. Figure S4H, Suppl. Table T1).

Taken together, labeling the upper TMD appears to report both on conformational changes related to pore opening and those that already happen at moderate pH and are more likely associated with ECD protonation. Introducing the N414K reduces acute desensitization and SSD at the level of the pore, but the fluorescence changes related to moderate pH could still be recorded from regions well below the molecular clutch.

### Peptides still bind to SSD-impaired channels

Many of the most potent ASIC1a modulators not only affect activation but also have profound effects on SSD. Two of these modulators are the endogenous neuropeptide Big dynorphin and the tarantula peptide toxin PcTx1, which likely act through an overlapping binding site involving the acidic pocket (28, 36, 46). We wanted to investigate if these peptides still affect channels where desensitization was largely abolished through the N414K mutation.

Big dynorphin rescues currents from SSD and has been shown to promote acidosis-induced cell death in cultured neurons (16, 47). Intervening with the ASIC1a–Big dynorphin interaction might thus be of pharmaceutical relevance. In mASIC1a WT, pH 7.2 introduces almost complete SSD (relative response 0.173 ± 0.10, Figure 5A, top and B, left, Suppl. Table T3). Adding 1 μM Big dynorphin did not immediately affect currents during preconditioning at pH 7.2 but almost completely rescued subsequent activation (relative response 0.91 ± 0.10, Figure 5A, top and B, left). To test a possible impact of Big dynorphin on K105C/N414K, we had to use pH 6.6 to introduce at least some SSD (relative response 0.59 ± 0.19, Figure 5A, bottom and B, right, Suppl. Table T3). At this pH, channels already activate substantially (relative response 0.33 ± 0.10). The addition of 1 μM Big dynorphin decreased pH 6.6 activation (relative response 0.19 ± 0.10) and showed some current rescue from SSD, albeit to a lesser extent than in the K105C mutant alone (relative response 0.79 ± 0.18, Figure 5A, bottom and B, right, Suppl. Table S3). VCF experiments on K105C/N414K revealed that applying Big dynorphin at pH 7.6 does not affect the current but introduces a slow-onset negative deflection of the fluorescence signal, indicating that the peptide binds to the channel. The introduced fluorescence change only fully returns to baseline once channels were activated with low pH. These results align with insights from other studies in K105C and WT channels, showing how Big dynorphin primarily stabilizes the closed and desensitized state but cannot bind the open state efficiently (16, 36, 46).

**Figure 5:**
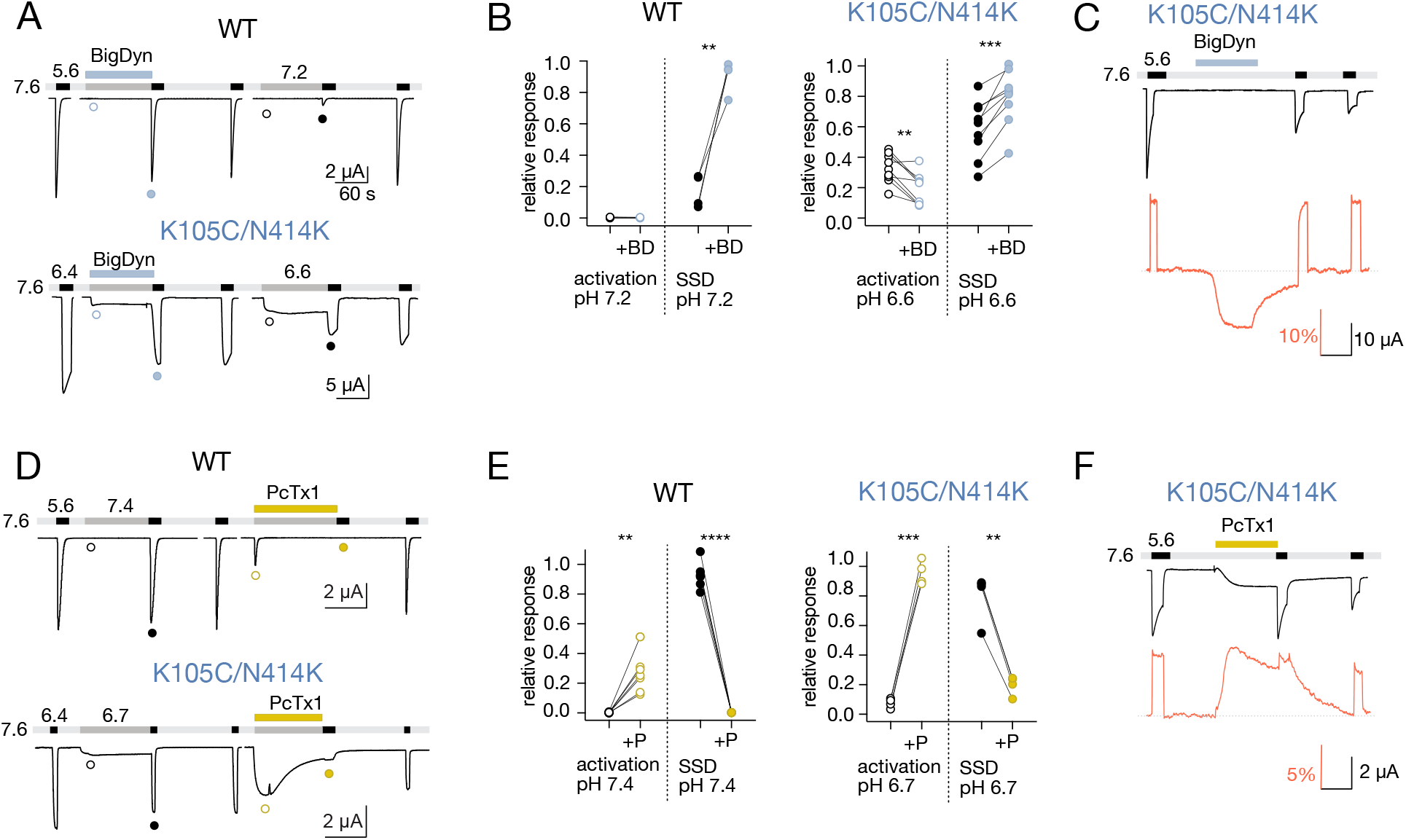
Effect of known peptide modulators on K105C/N4141K. **A)** Representative TEVC traces of WT (top) and K105C/N414K (boVom), showing the effect of 1 μM Big dynorphin (BigDyn) in an SSD protocol with a preconditioning pH of 7.2 or 6.6, respectively. **B)** Quantitative analysis of the data in panel A where the effect of Big dynorphin (+BD) is shown for both the activation with preconditioning pH (marked as an example in panel A with open circles), as well as the amount of SSD of the current (marked in panel A as closed circles). **C)** VCF trace with current in black and fluorescence in red, showing that BigDyn induces a slow-onset conformational change that only fully reversed back to baseline upon channel activation. **D)** TEVC traces of WT (top) showing how 30 nM PcTx1 at pH 7.4 can inhibit subsequent channel activation. To assess the effect on K105C/N414K (boVom), 100 nM PcTx1 at pH 6.7 was used in experiments similar to those for WT. **E)** Quantitative analysis as in panel B but for PcTx1 (+P). F) VCF recording of K105C/N414K showing the distinct effect of 300 nM PcTx1 on current (black) and fluorescence (red). All traces are on the same time scale. Statistical analysis: paired t-test ^**^=p<0.01, ^***^=p<0.001, ^****^=p<0.0001.

To investigate if we can reintroduce desensitization into K105C/N414K, we turned to the tarantula toxin PcTx1 (48). PcTx1 has been shown to reduce infarct volumes and have a neuroprotective effect in mouse models of stroke (8). PcTx1 potently shifts pH dependence of activation and SSD to lower proton concentrations, effectively introducing SSD already at pH 7.4 in WT channels (Figure 5 D, top and E, left, Suppl. Table T4) (49–51). Given the pH-dependent effect of PcTx1, we applied PcTx1 at 100 nM in pH 6.7 to the K105/N414K mutant and observed a distinct increase in channel activation during preconditioning (relative response 0.08 ± 0.03 (without) vs 0.96 ± 0.08 (with)), followed by enhanced SSD (relative response 0.80 ± 0.17 (without) vs 0.20 ± 0.07 (with), Figure 5D, bottom and E, right, Suppl. Table T4). In VCF experiments, PcTx1 also introduced a prominent conformational change similar to those previously seen in K105C (Figure 5F). These results clearly indicate that introducing the N414K mutation relatively stabilizes the open state of the channel, but PcTx1 can reintroduce desensitization.

Taken together, even channels that show reduced desensitization due to the introduction of N414K were still affected by peptide modulators known to affect mASIC1a desensitization, provided pH conditions were adjusted based on the pH sensitivity of the mutant.

## Discussion

In this study, we investigated the functional and conformational consequences of the mASIC1a β11-12 linker mutation N414K on acute desensitization, SSD, and tachyphylaxis. We found that the mutation retained pH-dependent conformational changes of the ECD and upper TMD. Functionally, the mutation impaired acute desensitization and SSD and introduced pronounced tachyphylaxis. Application of PcTx1, a modulator that enhances SSD in WT, partially restored SSD in the N414K mutant. Collectively, our work corroborates and expands our understanding of the molecular mechanisms involved in ASIC desensitization and emphasizes the important role of the β11-12 linker and its intricate interplay with pH-dependent conformational changes of the ECD.

### The N414K mutation impacts acute desensitization, SSD, and tachyphylaxis

Evidence from structural, functional, and computational experiments supports the role of the β11-12 linker as a molecular clutch that drives acute desensitization along with its surrounding region (12, 20, 29, 31, 32). Specifically, swiveling of chicken ASIC1 residues L414 and N415 appear to be at the heart of the molecular mechanism that controls the entry into and recovery from acute desensitization (20, 31). Here, the introduction of the mASIC1a N414K mutation reduced, but not abolished, acute desensitization in all four mutational backgrounds tested (mASIC1a K105C, V80C, E425C, and C70S/H72C) and the related rASIC3 isoform. While cysteine labeling itself also reduced acute desensitization, especially in positions K105C and E425C (Figure 1B, 2B, and 4B, Suppl. Figures S2A, S4B), all of the cysteine mutants showed almost complete SSD at preconditioning pH 6.9 (Figure 2E, 3E, 4F, Suppl. Figure S4G). However, together with the N414K mutation, none of the double mutants showed more than 20% SSD at pH 6.9 (Figure 2E, 3E, 4I, Suppl. Figure S4H). These data indicate that SSD is not abolished but rather strongly shifted to lower pH values and might remain incomplete even at low pH values or longer preconditioning times. This assumption is based on the observation that the pH-dependent activation is less affected by the N414K mutation than SSD, leading to a notable overlap between activation and SSD curves and thus generating currents that are somewhat similar to window-currents in ASIC3 (52) (e.g. Figure 2D and E). Precise assessment of the pH-dependent activation and SSD responses is not trivial in these mutants due to strong tachyphylaxis. For analysis, we normalized the size of the current of interest to the mean size of the two flanking low pH activations. This approach is not ideal because it assumes a linear decrease in current between the flanking low pH activations, although recurrent activations with the same pH show a non-linear decrease (e.g. Figure 2C). Yet, it appeared to be the most feasible option and has been successfully employed by others (e.g. (53)). TEVC data of the N414K equivalent position in human ASIC1a showed a similar decrease in acute desensitization and SSD (29), and patch-clamp data in chicken ASIC1 also reported markedly reduced desensitization kinetics in this mutant. The effect of N414K thus seems to be conserved across different species and methodological approaches, emphasizing its importance for acute desensitization and SSD.

Additionally, the introduction of the N414K mutation also led to pronounced tachyphylaxis in all backgrounds tested, including ASIC3, albeit to a much smaller extent. The presence and extent of tachyphylaxis have been shown to be highly dependent on species, isoform, and experimental setup and might be more pronounced in *Xenopus* oocyte recordings (20, 24, 25). Current evidence suggests that channels can only enter tachyphylaxis through the open state where permeating protons, and potentially also Ca^2+^ ions, govern the extent of tachyphylaxis (24, 25). The enhanced tachyphylaxis observed here is therefore likely not a direct mechanistic effect introduced by the N414K mutation but rather affects tachyphylaxis via stabilization of the open state, consequently enhancing the number of protons and Ca^2+^ ions flowing through ASIC1a. In line with this notion, we and others showed that tachyphylaxis increased with decreasing activation pH (Figure 1F). Factors that affect the resting state, such as washout time (Figure 1G) or washout pH (24) had little effect, although some recovery from tachyphylaxis was observed in patch-clamp recordings of rASIC1a (25). Activation time thus only affected the degree of desensitization in mutants that desensitize slowly and incompletely within the given activation time (Figure 1E, Suppl. Figure S1A, and B). In contrast, channels that desensitize within seconds showed the same extent of tachyphylaxis whether the channel was activated for 10 or 60 s (24). Permeating ions have been shown to change intracellular ion concentrations in patch-clamp recordings (54). Thus, local intracellular accumulation of permeating Na^+^ ions around ASIC1a could potentially lead to tachyphylaxis even in large cells such as oocytes if circumstances are extreme enough. In this case, we would expect longer washout times to alleviate some tachyphylaxis, either through dissipation of the altered local sodium gradient or reestablished sodium gradient due to the action of e.g. the Na^+^/K^+^ pump. However, this is clearly not what we observe experimentally (Figure 1G, Suppl. Figure S2D) and tachyphylaxis has been observed in the absence of extracellular sodium (24). In principle, differences in tachyphylaxis between ASIC3 and ASIC1a could arise from the higher Ca^2+^ permeability of ASIC1a (14), since increased Ca^2+^ permeability was linked to increased tachyphylaxis (24, 55). In the context of this study, we deem Ca^2+^ a rather unlikely major contributor since we did not add Ca^2+^ to the buffers. Rather, the N-terminal domain and permeating protons may play a role in tachyphylaxis (24). Yet, such a mechanism does not fully explain why the N414K/E425C mutant produced large non-desensitizing currents but shows less pronounced tachyphylaxis than other mutants (e.g. Figure 2B vs 3B), ultimately demanding a more in-depth study. Interestingly, over the course of multiple activations, acute desensitization decreased (e.g. Figure 1B and C). A possible explanation might be a shift in channel population where multiple activations lead to an accumulation of channels in a tachyphylaxis state, where the transition into an open state is slower than from the resting state (25). This would lead to less pronounced acute desensitization.

While we and others have shown a prominent role of the β11-12 linker in different types of desensitization, other parts of the protein also contribute to it. Here, the labeling position itself affected acute desensitization and SSD, and perturbations of desensitization can be observed from a wide array of mutations in the ECD (e.g. (40, 56), TMD (e.g. (57)) and the intracellular domain (e.g. (24)).

### Conformational changes in ECD and upper TMD are associated with desensitization

During activation, the conformational changes introduced by protonation of ECD residues propagate to the TMD and open the pore for ion flux. During acute desensitization, this coupling between ECD and TMD appears to be disturbed (13). Structural information of the open, desensitized, and resting state has shown that the upper ECD of the desensitized state matches that of the open state, while the lower ECD and TMD more closely resemble that of the closed state (12, 13). In line with these findings, our observations from VCF data with fluorophores in the upper (K105C) or mid (V80C) ECD present conformational changes during acute desensitization and SSD. At maximal SSD, the fluorescence signal is comparable to that of the acutely desensitized channels, suggesting that these states are similar or identical. Since channel gating kinetics vastly exceed our perfusion speed, we cannot directly comment on transitions between these states or delineate kinetic models. But data from others indicate that during activation with low pH, labelling in these positions tracks the transition from the resting to the active state (19, 40, 58) and that conformational changes already appear at sub-activating pHs (see also e.g. Figure 2D and E, 3D and E). While non-apparent openings suggested by the linear gating model (14) could explain changes in fluorescence at moderate pH, we deem it more likely that these fluorescence changes are a result of protonation events in the ECD that primarily lead to SSD in a model supported by others (19, 20). Disruption of the swiveling motion of the β11-12 linker via the N4141K mutation thus limited acute desensitization and SSD, but the dynamic conformational changes of the upper and mid-ECD appeared largely retained (Figure 2E and 3E).

Further, application of e.g. pH 6.9 to the K105C/N414K mutant resulted in activating currents despite the disrupted mechanism of the β11-12 linker (Figure 2D), indicating that conformational changes related to activation not strictly rely on the engagement of the molecular clutch, in line with similar observation by others (31). Activation seems to thus require a coordinated protonation of specific residues, some of which could be lying close to the ECD/TMD interface (59). Our data on tachyphylaxis further emphasized this point, showing how the conformational changes related to opening remained intact even when the channels underwent substantial tachyphylaxis. These data further indicate that tachyphylaxis does not originate from channel endocytosis, in line with earlier experiments that observed stable surface expression during tachyphylaxis (24).

Following the line of interpretation that the N414K mutation impedes some of the canonical ECD conformational changes from reaching the pore, we would expect to see conformational movements related to activation rather than desensitization when moving beyond the β11-12 linker to the top of the TMD. Indeed, labeling the top of TM2 in position E425C led to an increase in the fluorescent signal that coincides with increased activation, and a decrease when activation was reduced by SSD (Figure 4E and F). Yet, there was no precise match between the pH dependence of the fluorescence signal and activation, and our data indicate that even labeling at this position picks up on some of the conformational changes expected to come from the protonation of the ECD. Interestingly, the introduction of the N414K mutation shifted channel activation to lower pH and almost completely abolished acute desensitization and SSD within the pH range and preconditioning time assessed (Figure 4H and I). Consequently, almost no functional changes of the pore of the N414K/E415C mutation were detected at moderate pHs. In parallel, the fluorescence signal lost the pH-correlation with channel opening, leaving fluorescence signals similar to those obtained from labeling the upper and mid-ECD. While it is possible that the top of the TMD still undergoes some conformational changes even when desensitization is disrupted, we also deem it possible that the fluorophore not only reports on direct conformational changes of the upper TMD but also nearby conformational changes of the ECD, especially given the labeling position and the flexibility and length of the linker between side chain and fluorophore. Indications that the fluorescence signal is still somewhat related to channel opening, come from the observation that the fluorescence decreases in parallel to tachyphylaxis (Figure 4C and 4G). Our findings from labeling the top of TM1 indicate similar results, but the spread in the data makes a more detailed interpretation challenging (Suppl. Figure S4).

### Peptide modulation of channels with impaired desensitization

The ASIC1a modulators Big dynorphin and PcTx1 shift SDD curves to higher or lower proton concentrations, respectively (16, 50). Big dynorphin completely rescued SSD-induced currents in WT, but had a comparably minor effect on the K105C/N414K mutant. To assess the effect of the modulator on different mutants, the pH at which the modulators are applied has to be adjusted to comparable SSD levels in each mutant. In the K105C/N414K mutant, we thus needed to apply Big dynorphin at pH 6.6 to assess its ability to rescue SSD-induced currents. At this pH we already observed substantial channel activation. This likely makes the interaction unfavorable, since Big dynorphin primarily binds to the closed/desensitized state, and opening of the channel leads to unbinding of BigDyn (Figure 5C) (36). At pH 7.6, VCF experiments reveal that the conformational changes of K105C/N414K introduced by Big dynorphin appeared comparable to those seen for the Big dynorphin sensitive-K105C mutant (Figure C) (36), indicating that the resting state of the mutant can still interact with Big dynorphin. PcTx1competes with Big dynorphin for the same binding site but shifts activation and steady-state desensitization of ASIC1a to higher pH values. In the slow-desensitizing K105C/N414K mutant, 100 nM PcTx-1 had overall the same effect when applied at pH 6.7. Earlier studies have reported that 5 nM PcTx1 at conditioning pH 7.4, 7.5, and 7.6 neither decreased nor increased SSD of another slow-desensitizing Q276 mutant in hASIC1a (29). Given that we can reintroduce SSD in the K105C/N414K mutant using PcTx1 indicates that the binding partner can bridge the disrupted β11-12 linker to reintroduce desensitization.

### Conclusions

Desensitization in ASIC1a is complex and influenced by a myriad of factors. Manipulations to many sites within ASIC1a affect desensitization, but the β11-12 linker appears as a particularly critical region. The β11-12 linker residue N414 is also highly conserved even among relatively distant ASIC-like sequences (31, 60) suggesting that desensitization, or related mechanisms, are physiologically important. While the precise role of tachyphylaxis in physiology is still debated ((61, 62), SSD occurs at pH values around 7.1, and acidosis-inducing conditions such as ischemia, inflammation, or the microenvironment of tumors might therefore directly affect SSD in ASICs (9, 63–65). The effects of extracellular conformational changes at moderate pH might not only be relevant in the context of channel activation, but also on downstream signaling cascades (66–69). Understanding the precise molecular underpinnings of the different types of desensitization as a basic physiological function of ASIC is therefore highly important and could influence the development of ASIC-specific dugs.

## Author contributions

SAH and SAP conceptualized the project. CMH, ABT, JD, and SAH performed experiments and analyzed data. SAH and SAP wrote the original draft, and CMH, ABT and JD reviewed and edited the draft. SAP and SAH acquired funding.

## Acknowledgements

We acknowledge funding from the Lundbeck Foundation (R303-2018-3030 to SAH and R313-2019-571 to SAP), the Brødrene Hartmanns Fond, and the European Union’s Horizon 2020 research and innovation program under the Marie Skłodowska-Curie grant agreement no. 834274 (to SAH). We thank Dr. Han Chow Chua for comments on the manuscript.

