## Supplemental Figures for "Dynamic conformational changes of acid-sensing ion channels in different desensitizing conditions"

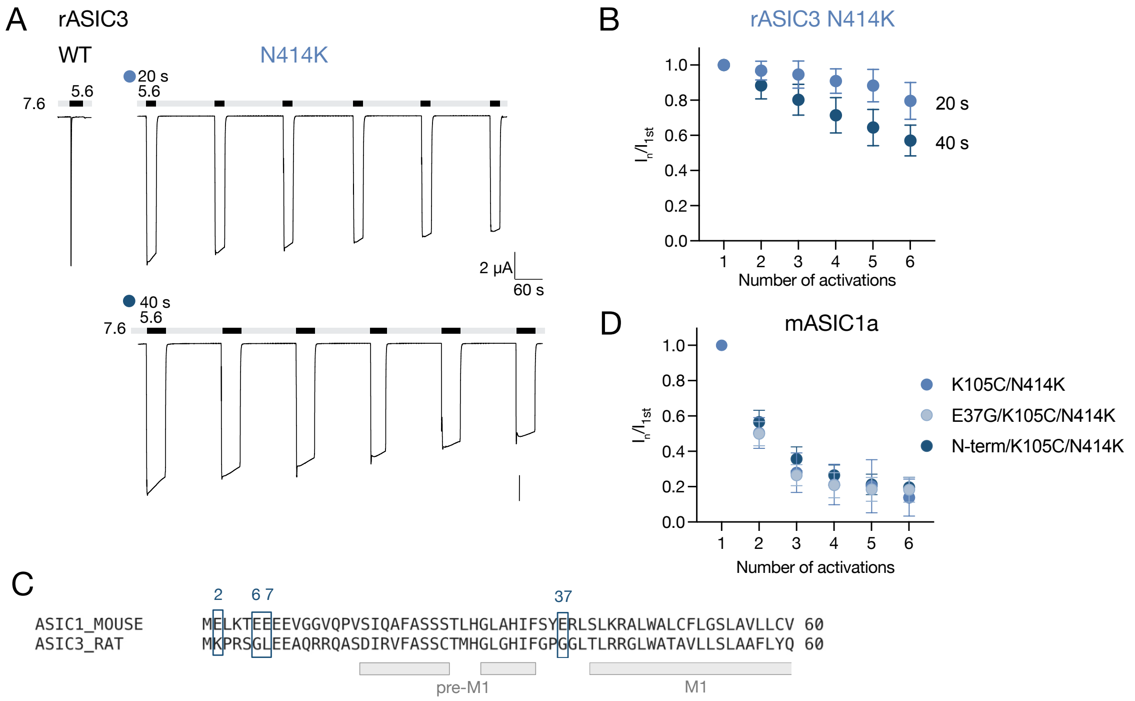


***Supplemental Figure S1: Effect of the N414K mutation in rASIC3 A)*** *Representative TEVC trace of rASIC3 WT (left) and mutation N414K (right) when activated with pH 5.6 for 20 s (top) or 40 s (bottom).* ***B)*** *Assessment of tachyphylaxis by comparing currents of recurrent activations to the current of the first activation (n=4-7).* ***C)*** *Sequence alignment of the N-terminal domain of mASIC1a and rASIC3, with residues that were mutated in boxes and numberd. Structural elements are shown below.* ***D)*** *Assesment of tachyphylaxis of N414K (also containing a K105C mutation) and mutants where either one (<u>E37G</u>/K105C/N414K) or all four (<u>N-term</u>/ K105C/N414K) highlighted glutamates shown in panel C were mutated to their conterpart in rASIC3 (n=4-8). All scale bars are 2 µA and traces are displayed on the same time scale. Data in graphs are presented as mean ± SD.*

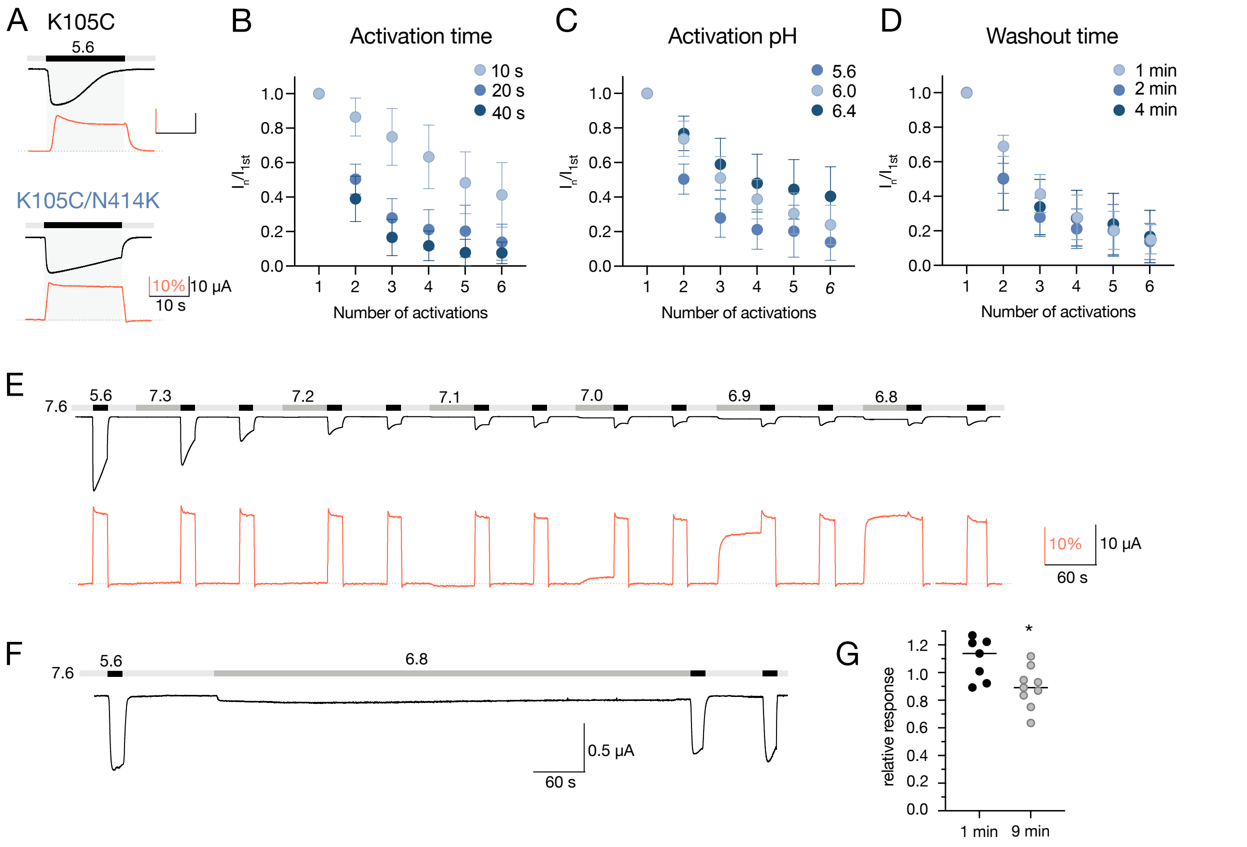


***Supplemental Figure S2: Effect of different activation protocols on tachyphylaxis in mASIC1a K105C/N414K. A)*** *Zoom-in on unfiltered VCF traces of K105C (top) and K105C/N414K (bottom) during activation with pH 5.5 with currents in black and fluorescence signal in red.* ***B)*** *Assessment of tachyphylaxis at various activation times by comparing currents of recurrent activations to the current of the first activation (n=4-9).* ***(C)*** *Same as in panel B but using different activation pHs (n=4-11).* ***D)*** *Same as in panel B but comparing tachyphylaxis using different washout times (n=4-8).****E)*** *Full VCF trace of the SSD recordings of K105C/N414K shown in Figure 2D, right.* ***F)*** *TEVC trace of an SSD recording of K105C/N414K, where channels were preconditioned with pH 6.8 for 9 min before subsequent activation.* ***G)*** *Comparison between activation after 1min and 9 min preconditioning in pH 6.8, respectively. Statistical analysis (non-parametric Welch's t-test). *=p<0.05. Data in B-D are presented as mean ± SD.*

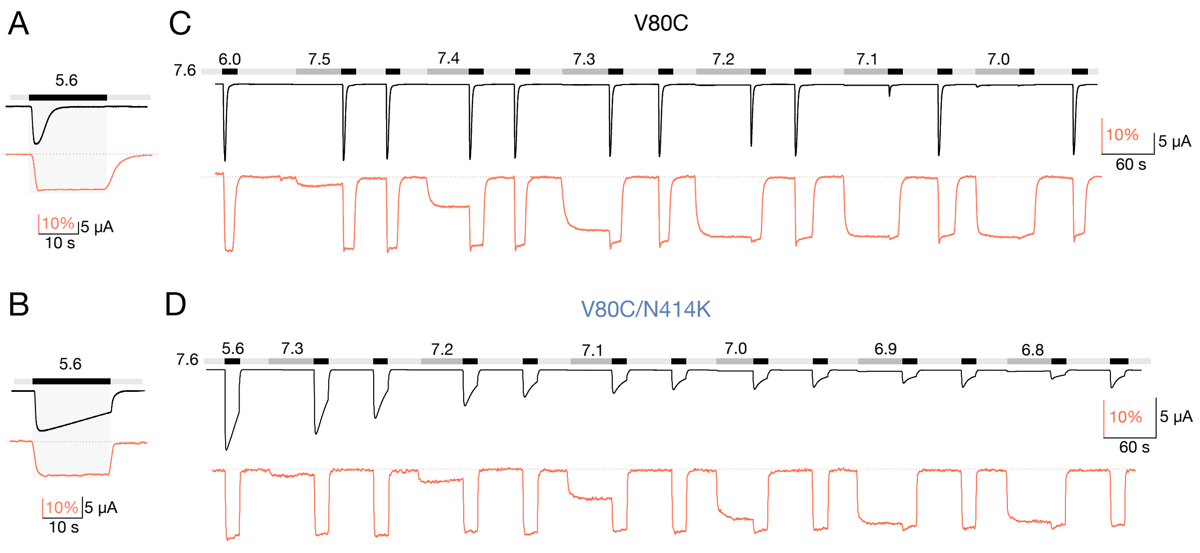


***Supplemental Figure S3: A)*** *Zoom-in on unfiltered VCF traces of V80C during activation with pH 5.5 with currents in black and fluorescence signal in red.* ***B)*** *Zoom-in on unfiltered VCF traces of V80C/N414K during activation with pH 5.5.* ***C)*** *Complete VCF traces assessing SSD in V80C.* ***D)*** *Complete VCF traces assessing SSD in V80C/N414K.*

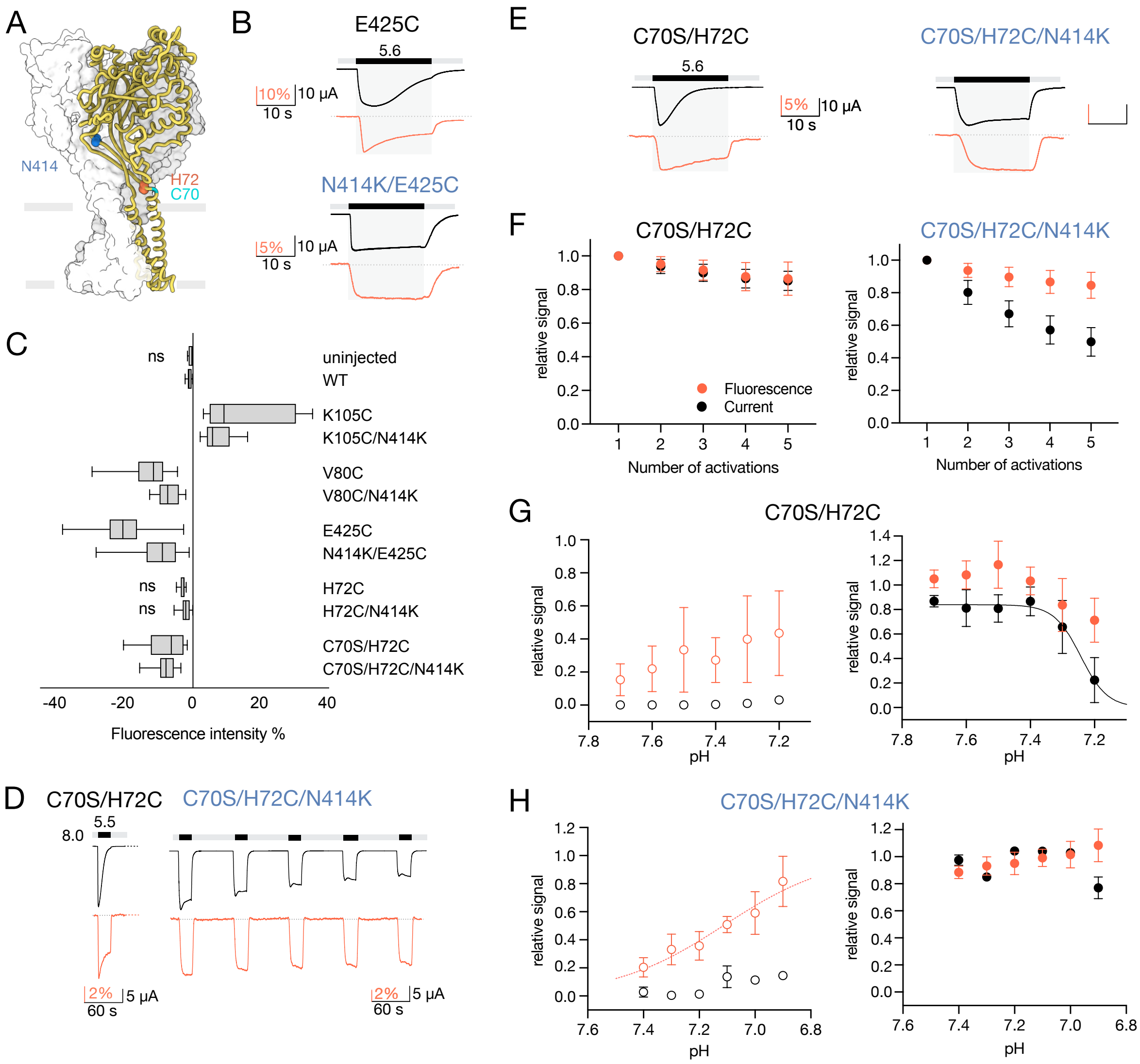


***Supplemental Figure S4: Conformational changes reported by labeling the top of TM1. A)*** *Structural overview of cASIC1 (PDB ID 6VTL) showing positions corresponding to mASIC1a N414 in blue, H72C used for fluorescent labeling in red, and C70 that was mutated to serine in cyan.* ***B)*** *Zoom-in on unfiltered VCF traces of E425C (top) and N414K/E425C (bottom) during activation with pH 5.5 with currents in black and fluorescence signal in red.* ***C)*** *Comparison of the fluorescence intensity recorded from different mutants. Besides uninjected oocytes, signals from H72C and H72C/N414K were the only ones that were not significantly different from WT.* ***D)*** *Representative VCF trace with the current in black and the fluorescence in red. The trace for C70S/H72C/N414K shows repeated 20 s activation with pH 5.6 with 1 min recovery in pH 7.6.* ***E)*** *Zoom-in on unfiltered VCF traces of C70S/H72C (left) and C70S/H72C/N414K (right) during activation with pH 5.5.* ***F)*** *Assessment of tachyphylaxis of C70S/H72C (left) and C70S/H72C/N414K (right) by comparing currents and fluorescence of recurrent activations to values of the first activation (n=12-16).* ***G)*** *Quantitative analysis of current and fluorescence evoked by moderate pH (left) and when evoked by low pH after preconditioning in moderate pH (right) as shown in Figure 4D-F (n=4-6).* ***H)*** *Same as in G, but for C70S/H72C/N414K (n=3-6). Data in graphs are presented as mean ± SD. Fluorescence intensity was compared to WT using an ordinary one-way ANOVA, ns = not significant.*
