## Supplemental Tables for "Dynamic conformational changes of acid-sensing ion channels in different desensitizing conditions"

**Supplemental Table T1:** Data summary of $pH_{50}$ values for the different mutants (mean, ±SD, n). Values without SD indicate that non-linear curve-fitting was not applicable within the pH range tested. Statistical analysis (unpaired Welch's t-test) between $pH_{50\_Fluorescence}$ and $pH_{50\_SSD}$ where appropriate, significant levels: **=p<0.01, ***=p<0.001; ****=p<0.0001

| Variant | $pH_{50\_Activation}$ | $pH_{50\_SSD}$ | $pH_{50\_Fluorescence}$ |
|---|---|---|---|
| K105C | 6.61 ± 0.07 (n=5) | 7.04 ± 0.06 (n=9) | 7.06 ± 0.06 (n=7); ns |
| K105C/N414K | 6.49 ± 0.25 (n=9) | < 6.6 | 6.93 ± 0.02 (n=7) |
| V80C | 6.65 ± 0.22 (n=7) | 7.17 ± 0.03 (n=4) | 7.36 ± 0.04 (n=4); *** |
| V80C/N414K | < 6.6 | ~ 6.6 | 7.11 ± 0.03 (n=5) |
| E425C | 6.87 ± 0.10 (n=19) | 7.17 ± 0.13 (n=11) | Act: 7.06 ± 0.07 (n=10); ****<br>SSD: 7.29 ± 0.04 (n=11); ** |
| N414K/E425C | ~ 6.4 | - | Act: 7.22 ± 0.15 (n=20) |
| C70S/H72C | - | 7.25 ± 0.04 (n=6) | - |
| C70S/H72C/N414K | - | - | Act: 7.05 ± 0.15 (n=4) |

**Supplemental Table T2:** Data summary of pH 5.6 activation after preconditioning in pH 6.8 for 1 or 9 minutes (mean ± SD), responses relative to pH 5.6. Statistical analysis (paired t-test), significant levels: *=p<0.05.

| Variant | 1 min | 9 min |
|---|---|---|
| K105C/N414K | 1.09 ± 0.15 (n=7) | 0.89 ± 0.15(n=9); * |

**Supplemental Table T3:** Data summary of the effect of 1 µM big dynorphin on WT and the K105C/N414K mutant (mean ± SD), all responses relative to pH 5.6. Statistical analysis (paired t-test) between applications with and without the peptide, significant levels: **=p<0.01, ****=p<0.0001, ns=not significant.

| WT | - | + 1 µM BigDyn |
|---|---|---|
| Activation pH 7.2 | 0.003 ± 0.003 (n=4) | 0.003 ± 0.002 (n=4); ns |
| SSD after preconditioning in pH 7.2 | 0.17 ± 0.10 (n=4) | 0.91 ± 0.10 (n=4); ** |

| K105C/N414K | - | + 1 µM BigDyn |
|---|---|---|
| Activation pH 6.6 | 0.33 ± 0.10 (n=9) | 0.19 ± 0.10 (n=9); ** |
| SSD after preconditioning in pH 6.6 | 0.59 ± 0.19 (n=9) | 0.79 ± 0.18 (n=9); *** |

**Supplemental Table T4:** Data summary of the effect of 30 nM PcTx1 on WT and 100 nM PcTx1 on the K105C/N414K mutant (mean ± SD), all responses relative to pH 5.6 activation. Statistical analysis (paired t-test) between applications with and without the peptide, significant levels: **=p<0.01, ***=p<0.001, ****=p<0.0001.

| WT | - | + 30 nM PcTx1 |
|---|---|---|
| Activation pH 7.4 | 0.002 ± 0.003 (n=7) | 0.27 ± 0.13 (n=7); ** |
| SSD after preconditioning in pH 7.4 | 0.93 ± 0.09 (n=6) | 0.002 ± 0.001 (n=8); **** |

| K105C/N414K | - | + 100 nM PcTx1 |
|---|---|---|
| Activation pH 6.6 | 0.08 ± 0.03 (n=4) | 0.96 ± 0.08 (n=4); *** |
| SSD after preconditioning in pH 6.7 | 0.80 ± 0.17 (n=4) | 0.20 ± 0.07 (n=4); ** |
